# BERLIN: Basic Explorer for single-cell RNAseq analysis and cell Lineage Determination

**DOI:** 10.1101/2023.07.13.548919

**Authors:** Gabrielle Nobles, Alyssa Obermayer, Ching Yao Yang, Tomas Zelenka, Thanh N Nugyen, Zhihua Chen, Jose Conejo-Garcia, Dorina Avram, Xiaoqing Yu, Y Ann Chen, Timothy Shaw

## Abstract

Single-cell RNA sequencing has revolutionized the study of immuno-oncology, cancer biology, and developmental biology by enabling the joint characterization of gene expression and cellular heterogeneity in a single platform. As of July 2023, the Gene Expression Omnibus now contains over 4000 published single-cell data sets, providing an invaluable opportunity for reanalysis to identify new cell types or cellular states as well as their defining transcriptional programs. To facilitate the reprocessing of these public datasets, we have devised a single-cell RNA sequencing analysis framework for data retrieval, quality control, expression normalization, dimension reduction, cell clustering, and data integration. Additionally, we have developed a Shiny App visualization platform that enables the exploration of gene expression, cell type annotations, and cell lineages through a user interface. We performed a re-analysis of single-cell RNAseq data generated from acute myeloid leukemia and tumor-reactive lymphocytes and found our pipeline to faithfully recapitulated the cell type assignment as well as expected lineage trajectories. Altogether, we present BERLIN, a single-cell RNAseq analysis pipeline that facilitates the integration and public dissemination of results from the reanalysis.

## Introduction

Single-cell RNA sequencing (scRNA-seq) has emerged as a robust and powerful method for examining the complexities of cellular heterogeneity, genetic expression, and cell state trajectories (Liu & Trapnell 2016). The Gene Expression Omnibus now contains more than 4000 scRNAseq data series. The current challenge is developing an effective strategy to standardize the reprocessing and data analysis with considerations to quality filtering, normalization, cell type identification, and differentiated lineages. While there are several flavors of sequencing protocol, including 10x genomics, Smart-seq2, Drop-seq, CEL-seq2, and Microwell-seq, these single-cell sequencing approaches generally have a high dropout rate in gene quantification.

Thus, careful quality control and filtering of low-quality cell is necessary (Chen et al. 2019; Ilicic et al. 2016). The integration of data generated from these different protocols also present unique challenges to the analysis due the variable capture efficiency and read coverage biases. One proposed strategy is to integrate the data by leveraging anchored gene features to correct for unwanted technical effects (Hafemeister & Satija 2019). Additional to addressing protocol variation, there is also a lack of standard in cell type assignment, which remains an open challenge in the scRNAseq analysis field (Lahnemann et al. 2020). Despite several automated cell type assignment algorithms are available, manual determination of the final cell type assignment is still necessary.

Here we generated BERLIN, a basic analytical pipeline protocol, that outlines a workflow for analyzing scRNAseq data. This protocol encompasses crucial steps, including quality control, normalization, data scaling, dimensionality reduction, clustering, and automated cell annotation. By following this protocol, users can effectively analyze scRNA-seq data, discern distinct cell populations, and gain valuable insights into cellular dynamics and functions. The results are explorable by ISCVA, an Interactive Single Cell Visual Analytics (Smalley et al. 2021) as well as with Shiny app, including UMAP, the EASY App, and the PATH-SURVEYOR Pathway Connectivity App. Overall, this protocol establishes a structured workflow for researchers, and a reproducible framework for scRNA-seq analysis and visualization.

## Methodology

### Installation Dependencies

BERLIN was developed with the open-source R programming language (v.4.2.2). The workflow leverages single cell RNAseq R packages that aid in clustering, cell annotation, and data manipulation. The single cell analysis is performed with Seurat (v.4.3.0) and Seurat helper packages, such as SeuratDisk (v.0.0.0.9020), and SingleCellExperiment (v.1.28.0). Cell annotation was performed with the R packages celldex (v.1.8.0), DoubletFinder (v.2.0.3), and SingleR (v.2.0.0) and for data cleaning and manipulation dplyr (v.1.1.2) and tibble (v.3.2.1) were used. The results of the BERLIN workflow can be visualized in the DRPPM-EASY (ExprAnalysisShinY), Shiny-UMAP, and PATH-SURVEYOR Pathway Connectivity applications that are compatible with R (v.4.2.2). There are package installation scripts for the BERLIN workflow and R Shiny applications through the GitHub page.

### Single-cell RNAseq Analysis Protocol

To maintain consistent and reliable cell annotation quality, the following steps were implemented to re-annotate cell types:

1. Unsupervised Louvain clustering algorithm: Utilized a multi-resolution approach to perform clustering, which aids in identifying distinct cell populations within the dataset.

2. SingleR cell type prediction: Employed the SingleR algorithm with multiple reference panels to predict cell types.

3. Gene marker identification: Identified specific gene markers associated with each cell type.

4. Visualization of dimensional reduction, annotations, and expression features: Utilized techniques such as dimensionality reduction (e.g., t-SNE or UMAP) to visualize the data in lower-dimensional space. Cell type annotations and expression features were integrated into the visualization, allowing for a comprehensive exploration of the scRNA-seq results.

5. Comparison of gene markers with public gene markers using the Pathway Connectivity app: This facilitated the assessment of overlap and similarities between the discovered gene markers and existing knowledge using the Jaccard distance calculation.

### Input Format

The input file should be a raw count matrix of single cells, with each row representing a gene and each column representing a specific cell. The values within the matrix indicate the raw expression counts or read counts of genes in each cell. It is important to ensure that the input file is in a compatible format, such as Comma-Separated Values (CSV), tab-delimited text (TXT), or Tab-Separated Values (TSV), as these formats can be easily read and processed by R and the Seurat package. The package also accepts CellRanger input files: matrix (matrix.mtx), barcode (barcode.tsv), and features (features.tsv). Examples of the input file can be found in https://github.com/shawlab-moffitt/BERLIN/tree/main/2-Single_Cell_RNAseq_Pipeline/Input

### Standard Quality Control

Perform quality control (QC) steps to filter out low-quality cells and genes. The percentages of mitochondrial genes and ribosomal protein genes were estimated for each cell. While the filtering criteria might be dependent on the library preparation protocol, cells with high mitochondrial and ribosomal gene content, or cells with low number of detectable features are typically removed from analysis.

~~~
qc.seurat <- function(seurat, species, nFeature) {
 mt.pattern <- case_when(
 species == “Human” ∼ “^MT-”,
 species == “Mouse” ∼ “^mt-”,
 TRUE ∼ “^MT-”
)
ribo.pattern <- case_when(
 species == “Human” ∼ “^RP[LS]”,
 species == “Mouse” ∼ “^Rp[ls]”,
 TRUE ∼ “^RP[LS]”
)
# Calculate percentage of mitochondrial and ribosomal genes
seurat[[“percent.mt”]] <- PercentageFeatureSet(seurat, pattern = mt.pattern, assay = “RNA”)
seurat[[“percent.rp”]] <- PercentageFeatureSet(seurat, pattern = ribo.pattern, assay = “RNA”)
# Filter cells based on QC criteria seurat[, seurat[[“percent.mt”]] <= 20 & seurat[[“nFeature_RNA”]] >= nFeature]
}
seurat_obj <- qc.seurat(seurat_obj, “Human”, 500)
~~~

### Expression Normalization

Unique molecular identifiers (UMI) counts are adjusted for library size and sequencing depth by dividing each UMI count by the total UMI count of that gene across all cells. This will ensure that expression values can be compared accurately. In Seurat, global-scaling normalization method “LogNormalize” that normalizes the gene expression measurements for each cell by the total expression, multiplies this by a scale factor (10,000 by default), and log-transforms the result.

This enables the preservation of the relative gene expression difference among cells.

~~~
seurat_obj <-NormalizeData(seurat_obj, normalization.method = “LogNormalize”, scale.factor = 10000)
~~~

### Generating Cell Cycle Scores

CellCycleScoring (Hao et al. 2021) function was used to determine the cell cycle activity for each cell. Specifically, the expression levels of certain genes associated with the cell cycle are analyzed and cell cycle scores are assigned to individual cells.

~~~
seurat_obj <-CellCycleScoring(object = seurat_obj, g2m.features = cc.genes$g2m.genes, s.features = cc.genes$s.genes)
~~~

### Scaling Data

Scaling normalizes and transform gene expression values, which helps remove unwanted technical variation, can improve downstream analyses. The scaling method used by default is the “LogNormalize” method, which performs a natural logarithm transformation followed by centering and scaling of the gene expression values. During the scaling process, the variables suspected to contribute to the batch variation can be specified in ‘vars.to.regress’ (nFeature_RNA and percent.mt in this case) are then regressed out in the analysis.

~~~
seurat_obj <-ScaleData(seurat_obj, vars.to.regress = c(“nFeature_RNA”, “percent.mt”), verbose = FALSE)
~~~

### Principal Component Analysis (PCA)

Conduct PCA on the scaled data to reduce the dimensionality of the dataset while preserving the most significant sources of variation. This step helps identify major sources of heterogeneity within the dataset.

~~~
seurat_obj <-RunPCA(seurat_obj, npcs = 30, verbose = FALSE, seed.use = 42)
~~~

### Nearest Neighbor and SNN clustering

Examining features derived by unsupervised selection as well as visualization of clusters in low dimensional space will reveal both biology as well as technical batch effects (Becht et al. 2018; Chen et al. 2019; Stuart et al. 2019). First, PCA is used to define cell neighbors in the low dimension space. Next, the algorithm performs clustering of the cells by constructing a shared nearest neighbor (SNN) graph, which connects cells based on their similarities in gene expression pattern. The Louvain algorithm is applied to optimize the modularity of a network by iteratively assigning nodes (cells) to different communities (clusters), which maximizes the within-community connections. Our pipeline performs the analysis at different resolutions to reveal high-and-low resolution groupings of cell types.

~~~
seurat_obj <- FindNeighbors(seurat_obj, reduction = “pca”, dims = 1:30)
seurat_obj <- FindClusters(seurat_obj, resolution = c(0.10, 0.15, 0.25,0.75))
~~~

### Doublet Finder

DoubletFinder is used to identify cells with putative doublets (McGinnis et al. 2019).

~~~
nExp <- round(ncol(seurat_obj) * 0.04) # expect 4% doublets
seurat_obj <- doubletFinder_v3(seurat_obj, pN = 0.25, pK = 0.09, nExp = nExp, PCs = 1:10)
~~~

### Automated Cell Annotation using SingleR and Celldex

SingleR and Celldex were applied to automatically annotate cell types. Celldex was utilized to download gene expression from the BlueprintEncodeData as part of the encode consortium, SingleR was used to compare and predict the cell type based on the gene expression of the reference dataset (Aran et al. 2019).

~~~
hpca.ref <- celldex::HumanPrimaryCellAtlasData()
dice.ref <- celldex::DatabaseImmuneCellExpressionData()
blueprint.ref <- celldex::BlueprintEncodeData()
monaco.ref <- celldex::MonacoImmuneData()
northern.ref <- celldex::NovershternHematopoieticData()
hpca.main <- SingleR(test = sce, assay.type.test = 1, ref = hpca.ref, labels = hpca.ref$label.main)
hpca.fine <- SingleR(test = sce, assay.type.test = 1, ref = hpca.ref, labels = hpca.ref$label.fine)
~~~

### Output files

Files that are generated from our pipeline include the following:

1. Metadata: Metadata is a data frame consisting of information derived from the pipeline:
  a. Cell Cycle Score (S.score,G2M.score, and Phase)
  b. QC percentage (percent.rp and perdent.mt)
  c. Doublets (pANN and DF.classfication)
  d. Resoulutions (integrated_snn_res or RNA_snn_res)
  e. Umap and Tsne coordinates
  f. Single R annotations
  g. Manually curated cell annotations
2. H5 Seurat-compliant file: Export the processed and analyzed data in the H5 Seurat file format. The H5 file is a hierarchical file that contains the expression values, dimensional reduction results, clustering information, metadata, raw counts and any other information stored within the Seurat object. The H5 Seurat object is the primary input file to the downstream post-processing.
3. H5 ISCVA-compliant file: The H5 ISCVA-compliant file is a specific file format designed to load and interact with the Interactive Single Cell Visual Analytics (ISCVA) application.

### scRNAseq post-Processing: Identifying defining markers

#### Input File

- H5 Seurat-compliant file: Load in the processed and analyzed data in the H5 Seurat file format. This file contains the expression values, dimensionality reduction results, clustering information, and metadata. It serves as a comprehensive representation of the analyzed single-cell RNAseq data.

### Post Processing: Find Markers

The H5-compliant Seurat object is loaded in using the LodH5Seurat() function from the SeuratDisk R package.

~~~
seurat_obj <- SeuratDisk::LoadH5Seurat(h5_file,
assays = c(“integrated”, “RNA”),
reductions = c(“pca”, “tsne”, “umap”, “rpca”),
graphs = NULL,
images = NULL,
meta.data = T,
commands = F,
verbose = F)
~~~

The gene markers for each Seurat cluster, with three or more cells, were derived using the FindMarkers() function from the Seurat package. From the markers identified, they were filtered into up-regulated (P<0.05 and LogFC>0) and down-regulated (P<0.05 and LogFC<0) datasets and written to an individual files for each cluster and a comprehensive excel notebook. This analysis can also be performed on an additional clustering criterion from the metadata file which the user can input in the beginning of the script. Furthermore, the upregulated gene markers from each cluster are used to develop a gene set of enriched markers per cluster, which is written out to a file for the user.

~~~
unfiltered_markers <- FindMarkers(object = seurat_obj, ident.1 = cells, min.pct = .25)
upreg_markers <- unfiltered_markers[unfiltered_markers$p_val_adj < 0.05 &
unfiltered_markers$avg_log2FC > 0, ]
dnreg_markers <- unfiltered_markers[unfiltered_markers$p_val_adj < 0.05 &
unfiltered_markers$avg_log2FC < 0, ]
~~~

### Post Processing: Subset 1000 cell matrix

To generate input files for the R Shiny Applications, the data is subset to a default 1000 cells by random sample with the ability for the user to set the seed. The assays of the Seurat object are iterated over and the subset count matrices, meta data, and H5 file are written to file.

### Output Files

1. Cluster Differentially Expressed Gene Markers: There are text files containing the DEG of each cluster as well as an excel workbook file containing all of the marker tables generated.
2. Cluster Labeled Gene Set: The gene markers from each cluster make up their indivdual geneset which is writted to a text and Rdata file.
3. Subset Scaled count matrix: The scaled count matrix has been normalized by log normalization and scaled and regressed based on ‘nFeature_RNA’ and mitochondria percentage.
4. Subset Raw count matrix: Raw count matrix contains the original counts.
5. Subset Normalized count matrix: The normalized count matrix is normalized by log normalization, but has not been scaled.
6. Subset Metadata file: A data frame that contains additional information associated with individual cells.

### Single-cell RNAseq Shiny App Explorer

We have developed a companion shiny app tool that provides users with the ability to visualize the output from BERLIN single-cell RNAseq analysis. This tool allows for the visualization of dimension reductions, major cell clustering at different resolution, and gene expression patterns. To demonstrate our tool’s functionality, we performed a scRNAseq reanalysis of a patient’s acute myeloid leukemia sample (van Galen et al. 2019) and CD69+103+ tumor-reactive lymphocytes derived from a ovarian cancer sample (Anadon et al. 2022). The Shiny app can facilitate cell type assignment using five different strategies: 1) cell type evaluation based on known gene markers. Using the AML scRNAseq data as an example, we first projected the cells in a Uniform Manifold Approximation and Projection (Figure 2A) unsupervised groupings defined based on the Louvaine clustering algorithm (Figure 2B). Lyz expression was high in cluster 4 and 7 (Figure 2C), which were annotated as Mono-like and GMP-like cells, respectively in the original manuscript (Figure 2D). The Lysozyme marker is a classical lineage-affiliated marker in myelomonocytic and granulocyte progenitor populations (Ye et al. 2003). 2) our pipeline provides a Jaccard pathway connectivity approach to compare gene markers derived from each cluster with publicly available gene markers. Through this approach, we validated that cluster 4 resemble mono-like cells and cluster 7 as GMP-like cells. We further identified cluster 3 as cDC-like cells and clusters 0 and 2 as HSC progenitors (Figure 3). 3) Gene markers can also be visualized as a barplots or boxplots for each cluster (Figure 4A), and predicted cell type annotation can be examined in each cluster (Figure 4B) and evaluated by a simple statistical test to evaluate the overrepresentation of a cell type in a cluster through a Fisher’s Exact Test (Figure 5). 4) our pipeline enables the real-time analysis of cell lineage by a Slingshot trajectory (Street et al. 2018), which can be jointly analyzed with the defined cluster and gene expression. We reanalyzed the T resident memory (TRM) cells derived from an ovarian cancer patient. Our trajectory analysis was able to identify four major cell types along the lineage trajectory consisting of TRM-stem, TRM-eff, TRM-prolif, and TRM-exhausted cells (Figure 6).

**Figure 1.**
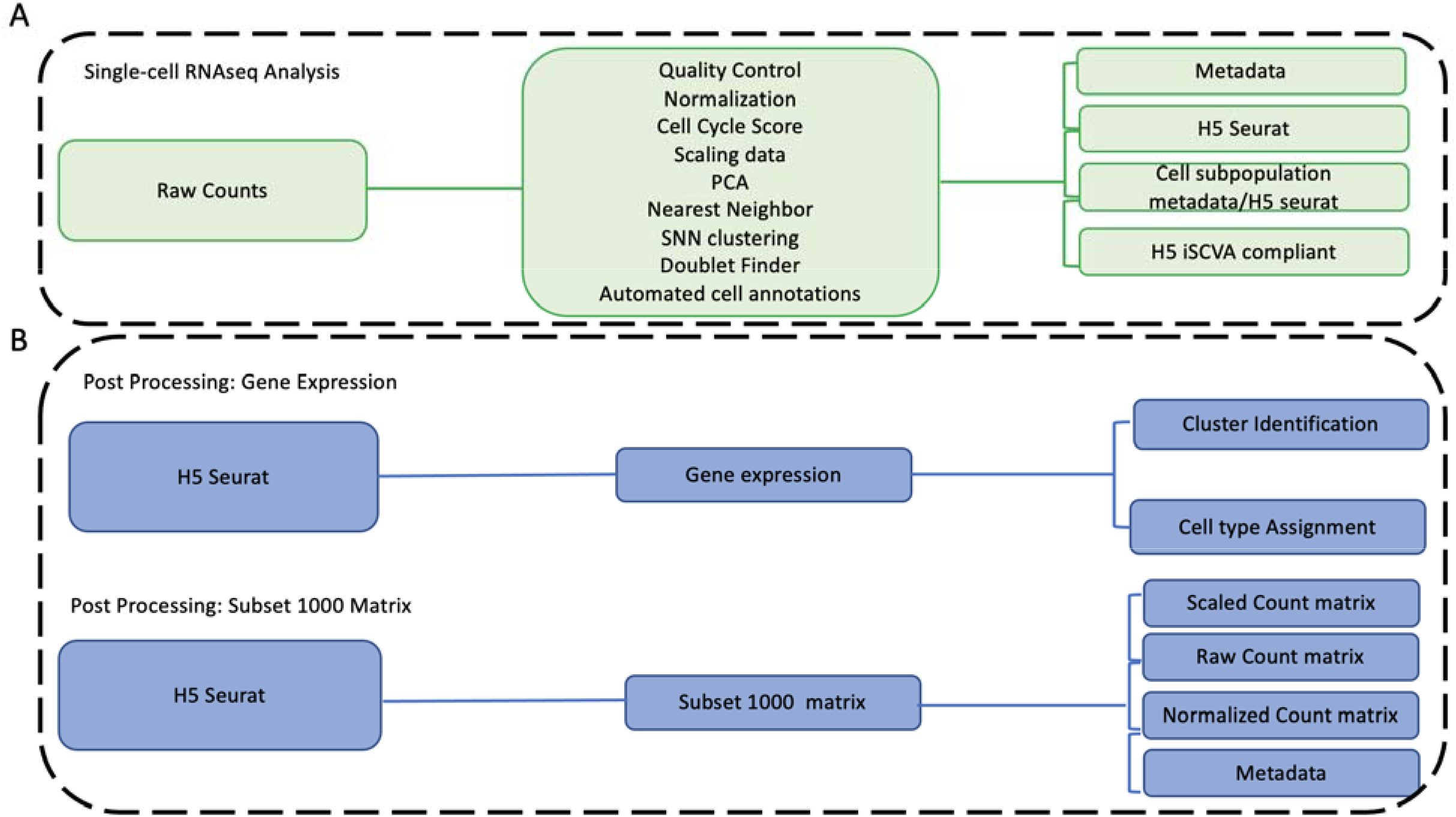
BERLIN single-cell analysis processing. A) Single-cell RNAseq analysis with standard quality control of barcode and gene features. Cells are annotated using SingleR backed by the CellDex database. B) Post processing of the single-cell analysis with H5 Seurat as the input. Unsupervised clustering is performed and cell types are assigned. Sampling of 1000 cells was performed to generate a smaller file for the Shiny app visualization.

**Figure 2.**
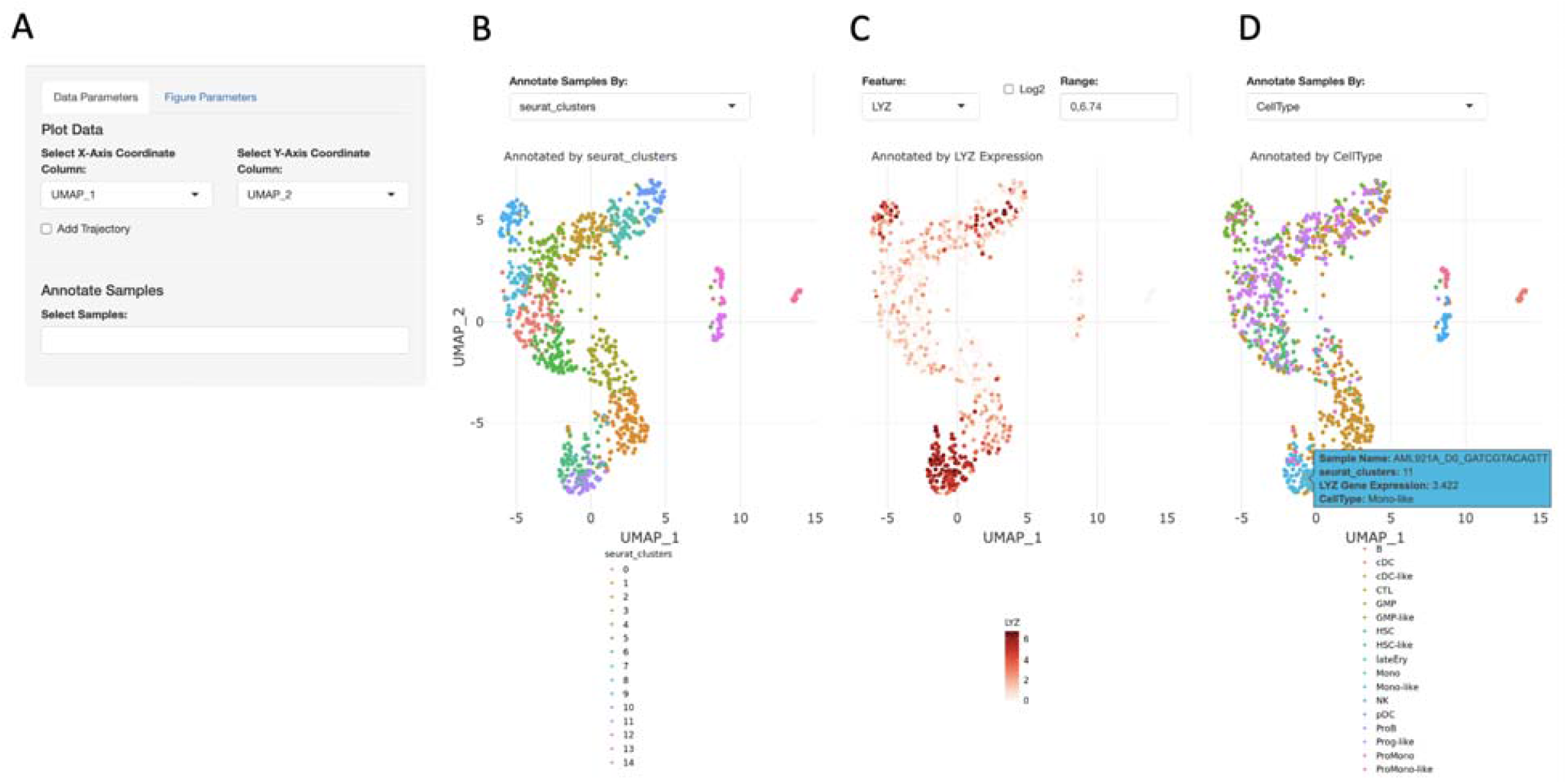
BERLIN-Shiny visualization of a scRNAseq data derived from an AML patient sample. Our app offers a multi-pronged visualization of both gene expression and cell type features. Dimension reduction was performed by the BERLIN pipeline and UMAP coordinates were selected for visualization **(A)**. Cells are colored based on the cluster defined by Seurat **(B)**. A subset of cells was demonstrated to be highly expressed with Lysozome (LYZ) **(C)**. The highly expressed cells were putatively annotated as monocyte-like **(D)**.

**Figure 3.**
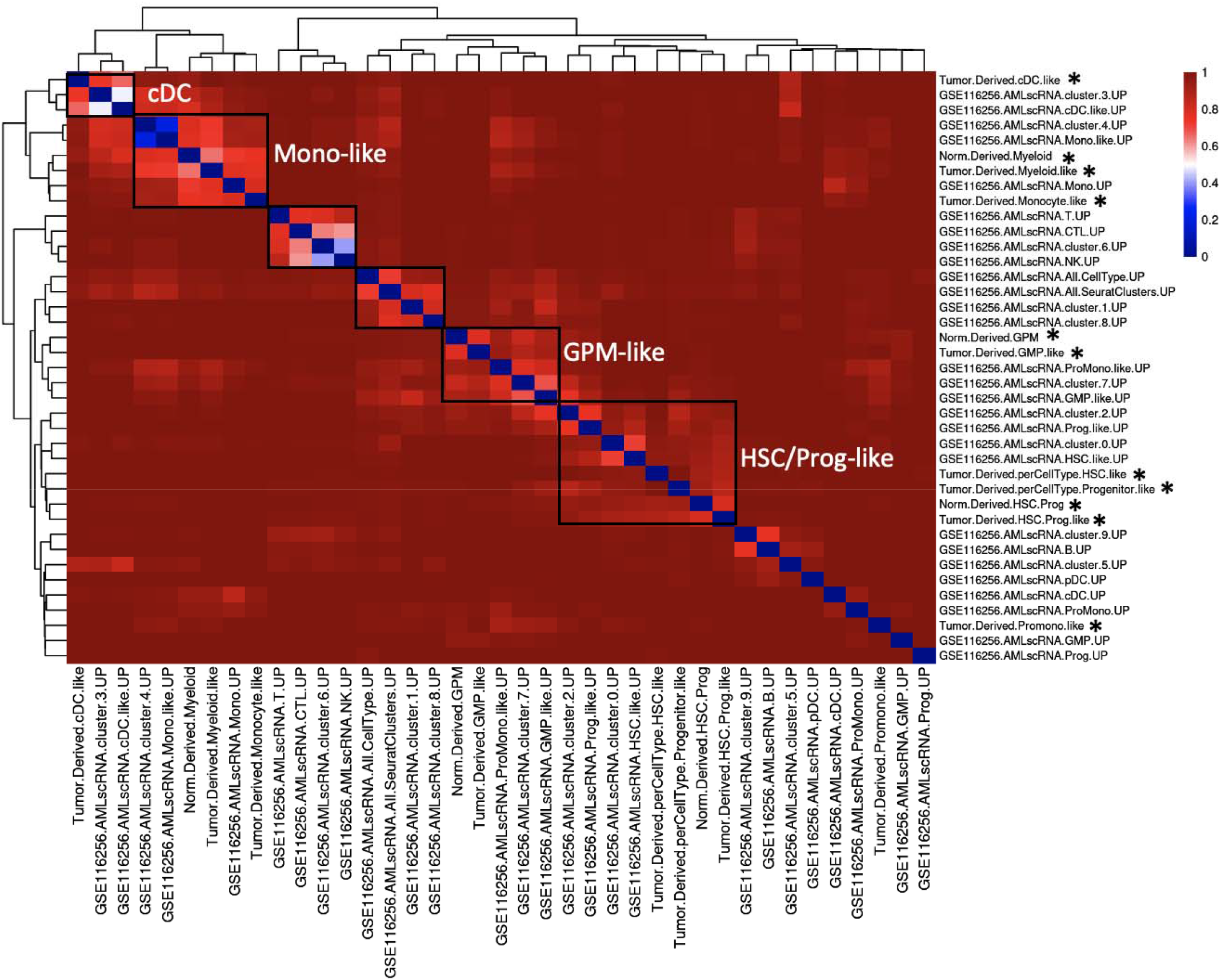
Cell type assignment with Jaccard connectivity analysis of AML cells. Upregulated gene marks in each cluster were compared to publicly available gene sets. As a separate validation, we also included up-regulated genes in each annotated cell type. The reference gene sets are marked with *

**Figure 4.**
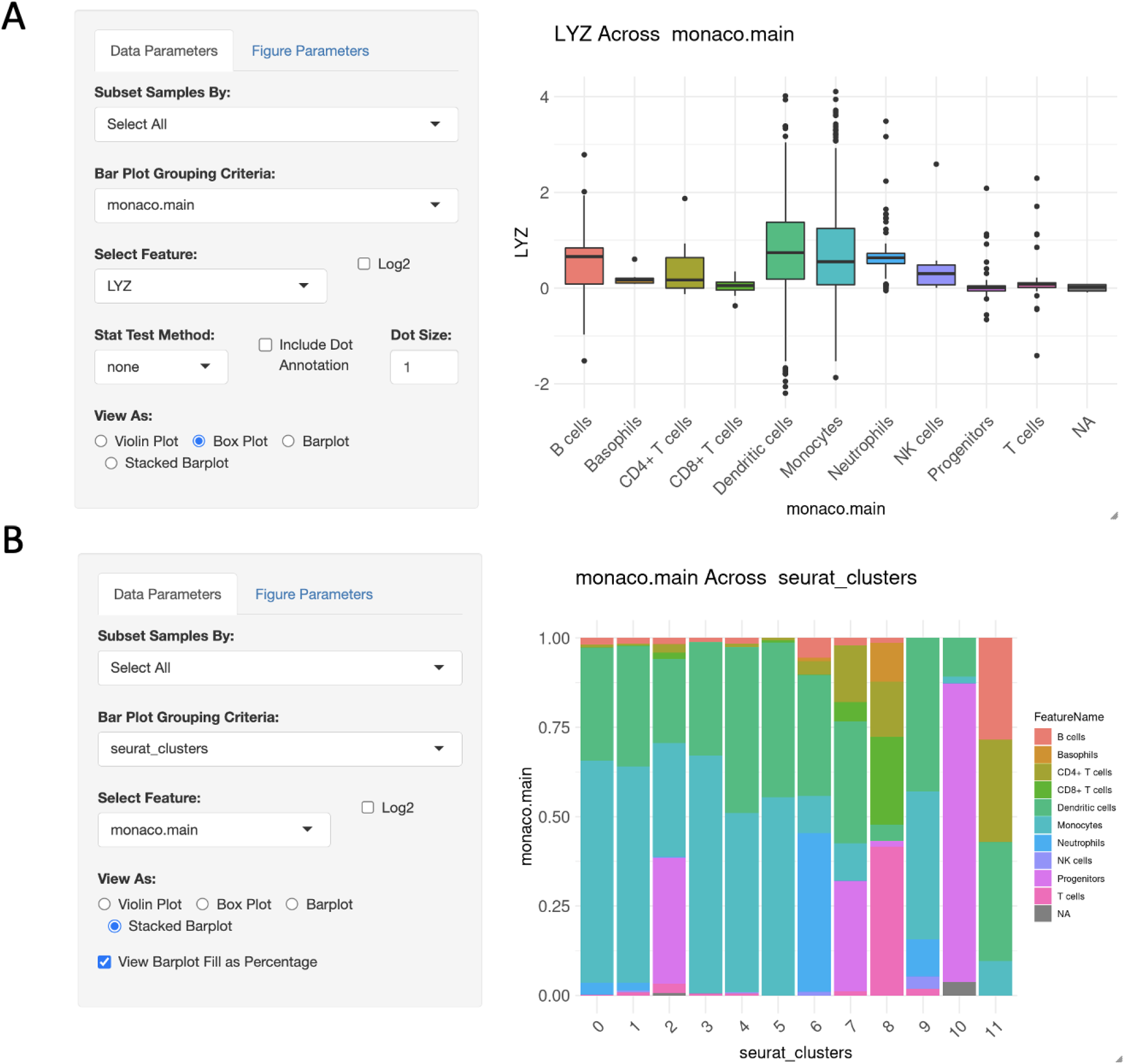
Feature comparison. A) Continuous features can be compared using violin or boxplot. B) Categorical features can be evaluated using stacked barplots.

**Figure 5.**
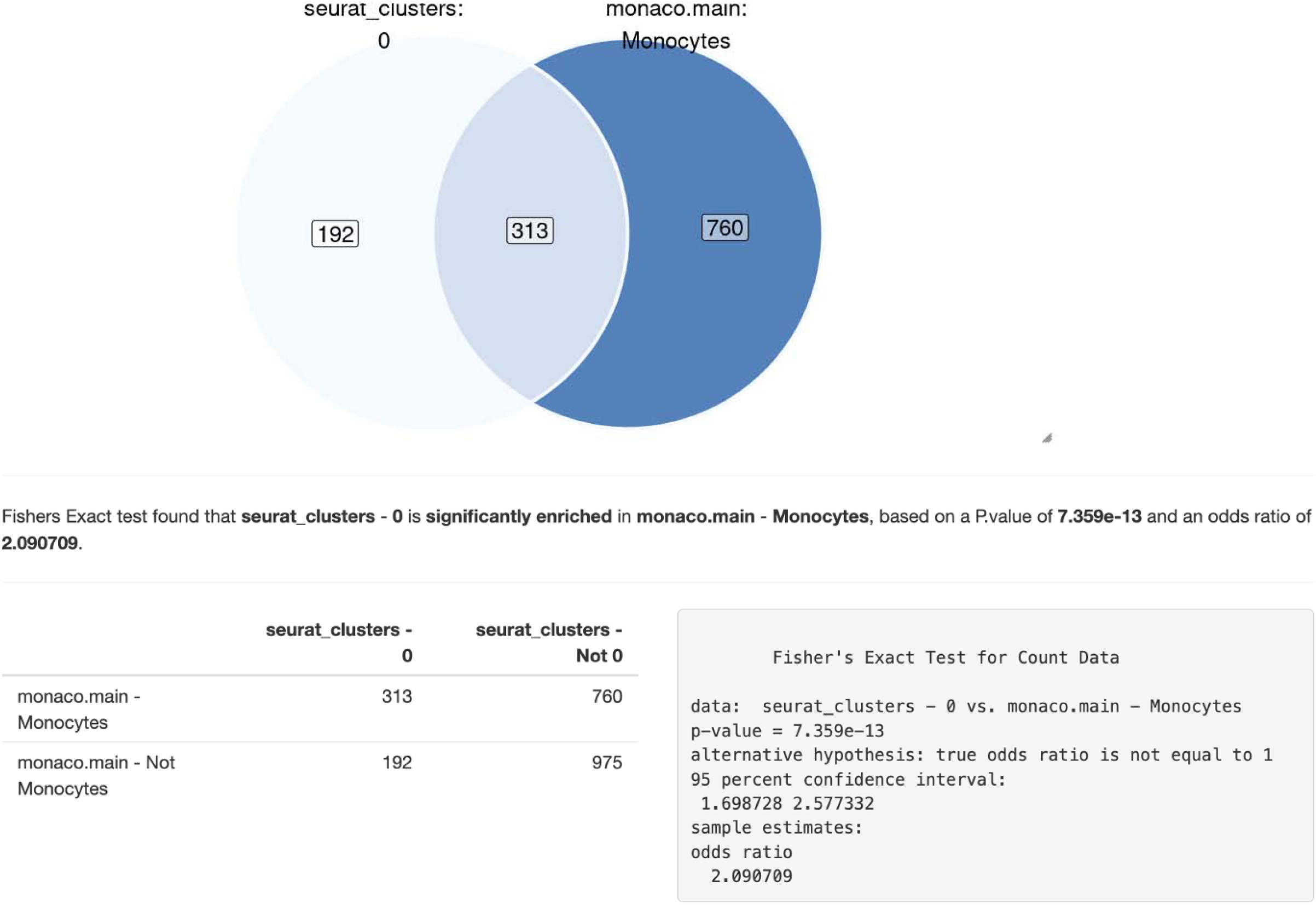
Enrichment of cell types or gene features. After generating a contingency table, enrichment can be performed using a Fisher’s Exact Test.

**Figure 6.**
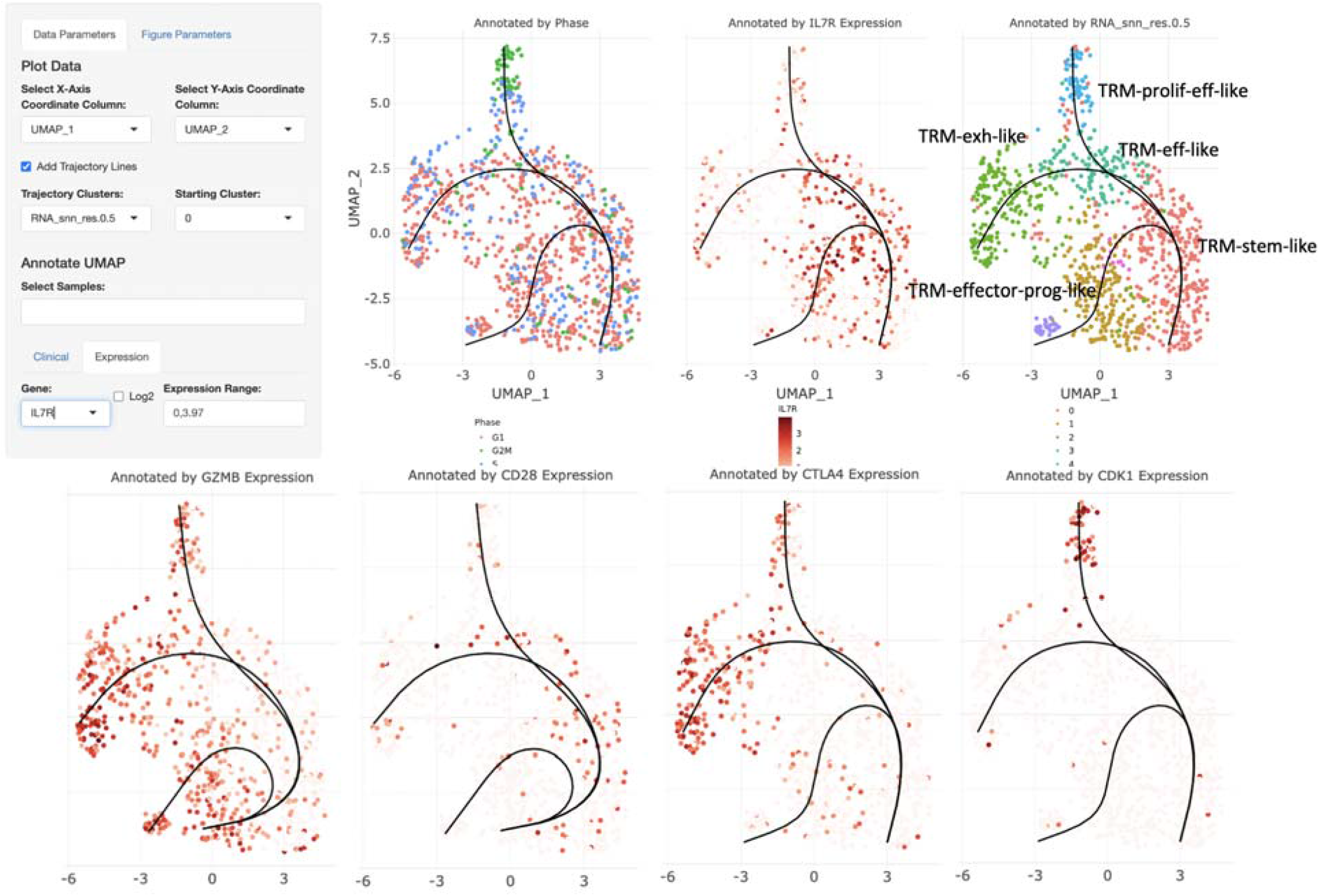
Single-cell lineage analysis of CD69+ 103+ tumor infiltrative T cells derived from ovarian cancer. A curve representing the cell differentiation trajectory is generated by Slingshot. The user can specify the cell grouping and the starting cluster of the analysis. The plot shows the differentiation trajectory of T resident memory cells supported by gene expression marker for stemness (IL7R and CD28), effector (GZMB), proliferation (CDK1), and exhaustion (CTLA4).

Interestingly, our analysis further revealed two distinct population of TRM-progenitor effector cells, which both contained high IL7R but with varying mid to high expression of GZMB (Figure 6). To facilitate an integrative gene expression analysis, samples can also be grouped based on genotype and explored by the DRPPM-EASY app (Obermayer et al. 2022). Finally, prognosis biomarkers derived from each cluster or cell types can also be evaluated using the PATH-SURVEYOR suite (Obermayer et al. 2023).

## Discussion

We have developed BERLIN, a basic unified protocol for single-cell RNAseq data analysis. With this pipeline, users can streamline analysis of publicly available scRNAseq datasets. As an interactive visualization tool is needed for a multidisciplinary research team to evaluate the same data set together, we further developed a Shiny app as a companion visualization tool.

The app can be effortlessly set up and applied to multiple single-cell datasets. The shiny app allows for in-depth analysis of cell lineage, gene markers, and cell differentiation trajectory.

Gene markers derived from our analysis can be further evaluated as biomarkers and associated with patient’s outcome through the PATH-SURVEYOR suite, and pathways enriched in each cell type can be examined through the DRPPM-EASY app suite. Collectively, these integrative applications will propel the user to gain additional insights into the gene regulatory mechanisms as well as the cell type’s potential as a prognostic biomarker.

In summary, our shiny app tool, coupled with the BERLIN pipeline, serves as a valuable resource for researchers in the fields of immuno-oncology, cancer biology, and developmental biology. Our tool will empower researchers to effectively analyze and virualize their single-cell data analysis, facilitating a deeper understanding of cellular dynamics and molecular processes in various domains of biological research.

## Data Accessibility

https://github.com/shawlab-moffitt/BERLIN/

## Example Shiny app is available

http://shawlab.science/shiny/BERLIN_GSE192780_TRM_UMAP_App/

